# Addition of 2′, 3′ *cis*-dialdehydes, 2′, 3′ *cis*-diols and phosphoryl groups to the 3′ end of oligonucleotides using periodate-oxidized nucleoside triphosphates and terminal deoxynucleotidyl transferase

**DOI:** 10.64898/2026.08.26.747364

**Authors:** Ross S. Anderson, Kenneth L. Beattie

**Affiliations:** Department of Biochemistry Baylor College of Medicine, One Baylor Plaza, Houston, TX 77030

**Keywords:** Terminal deoxynucleotidyl transferase, TdT, periodate-oxidized nucleotides, *cis*-dialdehyde, *cis*-diol, DNA 3′ phosphoryl group, affinity labelling

## Abstract

We present a simple and efficient way to add *cis*-dialdehydes, phosphoryl groups, or *cis*-diols to the 3′ end of oligonucleotides using periodate-oxidized nucleotides (o-NTPs) and terminal deoxynucleotidyl transferase (TdT). The 3′ *cis*-dialdehyde-modified oligos are generated by incubating TdT with an oligo for several minutes followed by addition of a o-NTP and incubated at 30°C for 30 minutes to an hour. After allowing the addition of the *cis*-dialdehydes, heating the reaction mixture at 90 – 95°C for 10 minutes yields oligonucleotides with 3′-phosphoryl groups. The 3′ *cis*-diol-modified oligos are synthesized by starting with 2′, 3′ *cis*-diol nucleotides (HO-NTPs).

The *cis*-dialdehyde modified oligonucleotides and *cis*-diols may then be used for a variety of investigations such as studying the interaction of proteins with the 3′ end of DNA, or possibly RNA. As an example, we demonstrate the efficacy of using an oligonucleotide modified with o-GMP at the 3′ end as an affinity label for TdT and identified a peptide fragment that has been shown to contain two of three aspartate residues found to be in the TdT active site.

**Highlights:**

- Terminal transferase can utilize periodate-oxidized nucleotide analogs to modify the 3′ end of oligonucleotides
- TdT can add a 2′, 3′ *cis*-dialdehyde to the 3′ end of an oligonucleotide
- Oligonucleotides modified with periodate-oxidized nucleotides can be useful affinity labelling agents for DNA-binding proteins
- Heating the 2′, 3′ *cis*-dialdehyde modified oligonucleotide in the presence of amino groups yields an oligonucleotide with a 3′ phosphoryl group
- Oligonucleotides with a 2′, 3′ *cis*-diol at the 3′ end can be produced using TdT with NaBH_4_-reduced 2′, 3′ *cis*-dialdehydes

## Introduction

Terminal deoxynucleotidyl transferase (terminal transferase, TdT; EC 2.7.7.31) catalyzes the random addition of deoxynucleoside 5′ monophosphates (dNMPs) to the free 3′-OH of a DNA primer in a template-independent manner. *In vivo* terminal transferase has been shown to play an important role in the generation of antibody and T-cell receptor diversity by randomly adding dNMPs to joins during antibody and T cell receptor maturation in B and T cells, respectively.

There are several naturally occurring *in vivo* enzymatic reactions that generate RNA or DNA with a 3′ phosphoryl group. Examples are RNase I and RNase A ribozymes which generate cleavage products with 3′ phosphoryl groups [1]. DNase II generates DNA with a 3′ phosphoryl group in the degradation of DNA during apoptosis [1]. Certain antibacterial medicines are also known to generate damaged DNA with a 3′ phosphoryl group [1]. A common *in vitro* way to chemically generate RNAs with a 3′ phosphoryl group is by alkaline hydrolysis [2].

Terminal transferase is commonly used as a molecular biology tool in various studies involving tailing of DNA, examples are the adding of either homo- or heteropolymer tails to plasmids or cDNA for *in vitro* mutational studies, the synthesis of XNA oligonucleotides (xenonucleic acids, oligonucleotides containing nucleotides modified in various ways), producing synthetic homo- or heteropolymers of single-stranded DNA, labeling the 3′ ends of DNA with radioactive nucleotides, for RACE (Rapid Amplification of cDNA Ends), and TUNEL (Terminal deoxynucleotidyl transferase dUTP Nick End Labeling) assays in the detection apoptosis. For a more comprehensive review of properties and applications of TdT see Zhang et al. [3].

Here we present a simple and effective procedure whereby oligonucleotides may be modified at their 3′ ends by the addition of 2′, 3′ *cis*-dialdehydes, 2′, 3′ *cis*-diols or a 3′ phosphoryl group using periodate-oxidized nucleotides (o-NTPs) and terminal transferase. We also demonstrate the efficacy of using o-NMP-modified oligos as affinity labels for proteins which interact with the 3′ ends of nucleic acids.

## EXPERIMENTAL

### Materials and Methods

#### (a) Oligonucleotide (p(T)_15_) was purchased from Pharmacia

Non-radioactive nucleoside triphosphates (NTPs) were purchased as 100 mM solutions from New England Bio Labs. Radioactive nucleotides were obtained from ICN. Bovine terminal transferase preparation of was highly purified on a monoclonal antibody column by Dr. Fred Bollum’s group at Supertechs, Rockville, MD, and through United States Biochemical. The preparation consisted of the 44 kDa α, β-heterodimer (95 – 98%) and the 45 kDa monomer (2 – 5%) as judged by silver and Coomassie Blue staining of SDS-polyacrylamide gels and western blot analysis.

Calf intestine alkaline phosphatase and T_4_ polynucleotide kinase were purchased from New England Bio Labs.

Polyethyleneimine (PEI) cellulose F_254_ plates were purchased from Millipore. Sodium borohydride and sodium periodate were purchased from Sigma-Aldrich.

All oligonucleotide modifying reaction mixtures contained 50 mM HEPES, pH 7.8, 7 mM Mg^2+^, or 100 mM cacodylate, pH 7.2 and 2 mM Co^2+^, 0.5 mM o-NTP, 10 µM p(T)_15_, and 0.3 µM TdT (assuming an M_r_ = 44 kDa) in a final volume of 30 µL. Reactions were carried out at 30°C in sterile 12 x 75 mm disposable borosilicate tubes obtained from Fisher Scientific.

#### (b) Preparation of Radiolabeled Oligonucleotides and NTPs

Stock solution of p(T)_15_ was prepared by suspending the lyophilized oligonucleotide in 100 – 300 μL of 10 mM HEPES, pH 7.8. Radiolabeled oligonucleotides labeled on the 5′ end were prepared with [γ-^32^P]ATP and T_4_ polynucleotide kinase according to Sambrook et al. [2].

Stock solutions of nucleoside triphosphates (10 – 20 mM) with a radiospecific activity of 100 – 200 cpm/pmol were prepared by adding 500 μCi of [α-^32^P]NTP (radiospecific activity ∼ 3000 Ci/mmol) to 100 – 200 μL of a 100 mM NTP solution and bringing the final volume to 1 mL with 10 mM HEPES, pH 7.8.

#### (c) Synthesis of 2′, 3′-cis dialdehyde NTP (o-NTP)

In all experiments freshly prepared o-NTP was used as o-NTPs in older preparations have a tendency polymerize forming structures that may resemble the DNA substrate [4]. Periodate-oxidized NTPs were synthesized in our lab as described by Easterbrook-Smith et al. [5].

**Fig. 1.**
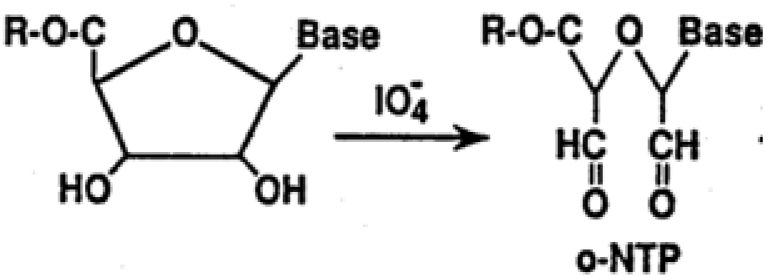
Periodate oxidation of the 2′ and 3′ carbons of the ribose moiety to aldehyde groups

Briefly, 25 – 50 μL of a 100 mM [α-^32^P]NTP solution was brought to 400 μL with sterile, distilled water. To this was added 200 μL of a freshly prepared 0.45 M solution of sodium periodate, pH 7.0. The solution was allowed to stand for 1.5 – 2 hours at 4°C in the dark. Progress of the reaction was monitored by spotting 1 or 2 μL onto polyethyleneimine (PEI) cellulose F_254_ thin-layer chromatography plates and developed in either 2 M LiCl [4] or 0.8 M (NH_4_)HCO_3_ [5]. Chromatographed compounds were located by UV-shadowing or autoradiography. Unreacted NTP migrated several centimeters from the solvent front and several centimeters from the origin, while o-NTP remained at the origin. The reaction was quenched by addition of 20 μL ethylene glycol. Periodate-oxidized NTP was purified from other reaction components by loading the entire reaction mixture onto a sterile Sephadex-G10 column equilibrated with sterile, distilled water at 4°C. Elution was with sterile water. Fractions of 1 mL were collected and assayed for the presence of iodate by a method described by Hinrichs and Eyzaguirre [6]. Only iodate-free fractions were pooled and lyophilized. The lyophilized materials were suspended in sterile water and the concentration measured spectrophotometrically at 252 nm with a Beckman DU70 spectrophotometer. The radiospecific activity of purified radiolabeled o-NTP was 200 – 1000 cpm/pmol.

#### (d) Synthesis of 2′, 3′-cis diol NTP (HO-NTP)

Reduced periodate-oxidized NTPs were synthesized as described by Lowe and Beechey ([4].

**Fig. 2.**
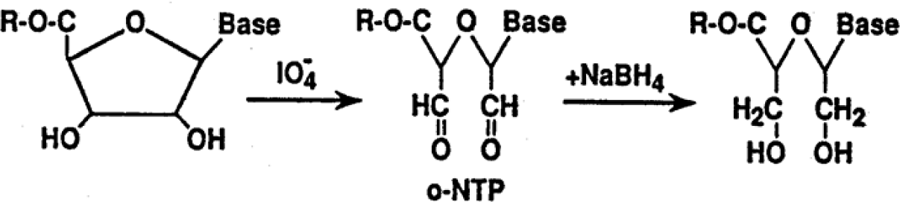
Synthesis of *2′, 3′-cis diol* NTP starting with o-NTP. The two aldehyde groups are reduced to alcohols

Briefly, a 4 M solution of NaBH_4_ was freshly prepared in 10 mM HEPES, pH 7.0. Aliquots of this solution were added to an o-NTP solution to a final NaBH_4_ concentration that was approximately a10-fold molar excess over the concentration of aldehyde groups. Only iodate-free fractions of o-NTP were used. The mixture was then incubated for 2 hours at 4°C. The reduction reaction was monitored by spotting 2 μL aliquots onto a PEI-cellulose F_254_ thin-layer chromatography plate and developed with 2 M LiCl or 0.8 M (NH_4_)HCO_3_. Spots were located by UV-shadowing, or autoradiography. The reaction was complete within two hours as judged by the disappearance of UV-absorbing material remaining at the origin and by the lack of detectable aldehyde groups when sprayed with 2,4-dinitrophenylhydrazine [4]. The reaction mixture was applied to a sterile Sephadex G-10 column and eluted with sterile water and fractions of 1 mL were collected. Fractions containing HO-NTP were located by thin-layer chromatography as described above. Appropriate fractions were pooled, lyophilized, and stored desiccated at -20°C. Immediately before use the lyophilized HO-NTP was suspended in sterile water and the concentration determined spectrophotometrically at 252 nm or 260 nm.

#### (e) Polyacrylamide gel electrophoresis

All PAGE analyses of modified DNA were done with a 12% non-denaturing polyacrylamide gel run at a constant 800V in a Tris-Borate-EDTA buffer (Bio Rad, Inc.).

Analysis of NTP and DNA protection against o-NTP modification were done with a 10 % resolving gel and a 4% stacking gel and run at a constant 150V. All SDS-PAGE analysis of modified protein or peptide fragments samples were incubated with NaBH_4_ for 30 minutes to 1 hour prior to loading on the gel.

#### (f) Peptide mapping by SDS-PAGE

Approximately 10 μg of TdT was incubated with 0.5 mM [α-^32^P]dGTP for 10 minutes at 30°C in activation buffer. In protection experiments TdT was incubated for 30 minutes with either 3 mM dGTP or 100 μM p(dT)_15_ prior to addition of o-GTP. Labeling took place in a 1.5 mL Eppendorf microcentrifuge tube. The reaction was terminated by the addition of NaBH_4_ to a final concentration of 34 mM and allowed to stand for 2 hours at room temperature. The reaction mixture was brought to 6% trichloroacetic acid by addition of 100 μL of 72% trichloroacetic acid [7–9]. Precipitation took place on ice for 30 minutes to 1 hour. The precipitate was pelleted by centrifugation for 30 minutes, then washed with 200 μL of cold HPLC-grade acetone three times, and air dried. After drying, 200 μL of 70% trifluoroacetic acid was added to the dried pellet and several crystals of cyanogen bromide (CNBr) were added and the tube sealed in the presence of argon. The reaction was allowed to proceed for 24 hours at room temperature in the dark. The trifluoroacetic acid and CNBr were then removed by speed-vac drying. The resulting peptide pellet was neutralized by the addition of 50 μL of ethanolamine, or 50 μL of 1 M (NH_4_)HCO_3_, which was removed by speed-vac drying. This step was then repeated. The final neutralized pellet was then subjected to SDS-PAGE.

Cyanogen bromide fragments were analyzed by SDS-PAGE as described by Schägger and von Jagow [10]. In these analyses a 16.5% total acrylamide (T), 3% crosslinker (C) separating gel was overlaid by a 2 cm layer of 10% T, 3% C gel which in turn was overlaid by a1 cm 4% T, 3% C stacking gel. Electrophoresis was carried out at 30 V until tracking dye entered the stacking gel, the voltage was then raised to a constant 85 V until the tracking dye ran off the bottom edge of the separating gel (approximately 18 hours). Myoglobin peptide standards (Sigma-Aldrich) were used to assess the relative sizes of the peptide fragments.

Resolved CNBr fragments were transblotted onto Immobilon-P (Millipore, Inc) in CAPS buffer, pH 11.0. Complete transfer was obtained within 2.5 hours at a constant 70 V at 4°C. Radio-labeled fragments were located by autoradiography and excised for sequencing. Sequencing was performed by Dr. Richard Cook, director of the protein core facility at Baylor College of Medicine.

### Results and Discussion

#### (a) Demonstration of 2′, 3′-cis dialdehyde NMP addition

Purified 5′ end-labeled p(dT)_15_ and unlabeled o-GTP were incubated with TdT and the fate of the oligonucleotide was monitored over time by non-denaturing PAGE (Fig. 3a). Lanes 1 through 9 represent reaction products over the course of 24 hours. Aliquots were not heated prior to loading on the gel. The fact that no other product bands formed above the upper bands indicates that the 3′ end of p(T)_15_ was modified such that no further polymerization could take place. An aliquot of the reaction from lane 9 (representing 24 hours) was subjected to TLC, developed with 2M LiCl, and subjected to autoradiography (Fig. 4). Consistent with the oligo possessing aldehyde groups at its 3′ end, the p(T)_15_ remained at the origin. The presence of unmodified p(dT)_15_ through 24 hours indicates that the DNA substrate was not limiting. The concentration of o-GTP in both experiments was 0.5 mM, the p(T)_15_ concentration was 10 µM, and the TdT concentration was 0.4 μM, therefore, all the DNA starting material should have been modified within the allotted time. Thus, TdT is inhibited by o-GTP, but not before undergoing several catalytic cycles. If each enzyme molecule underwent one catalytic cycle before being inhibited, then only a small fraction of the 5’ end-labeled p(dT)_15_ would have been converted to form product band.

**Fig. 3a.**
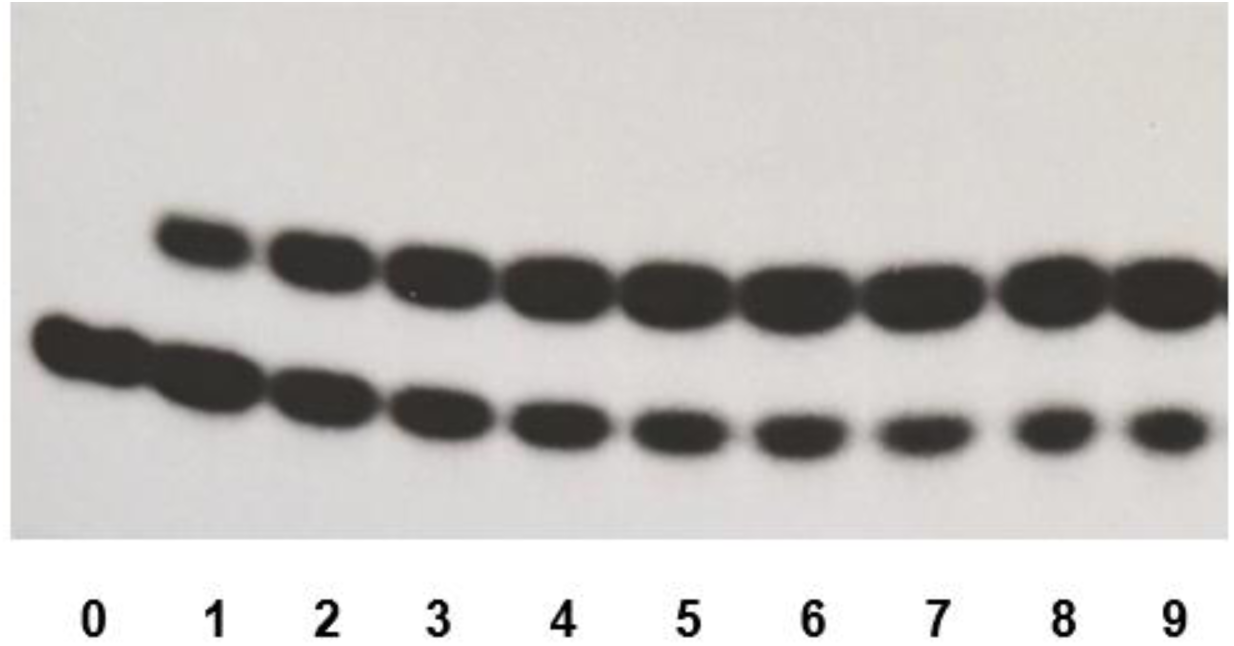
Modification of 5′ end-labelled p(T)_15_: **Lane 0:** 5′ end-labelled p(T)_15_, **Lanes 1 – 9:** 5′ end-labelled p(T)_15_ modified using o-GTP. At various times over 24 hrs. aliquots were removed and prepared for gel analysis. The enzyme was pre-incubated with the p(T)_15_ prior to o-GTP addition

**Fig. 3b.**
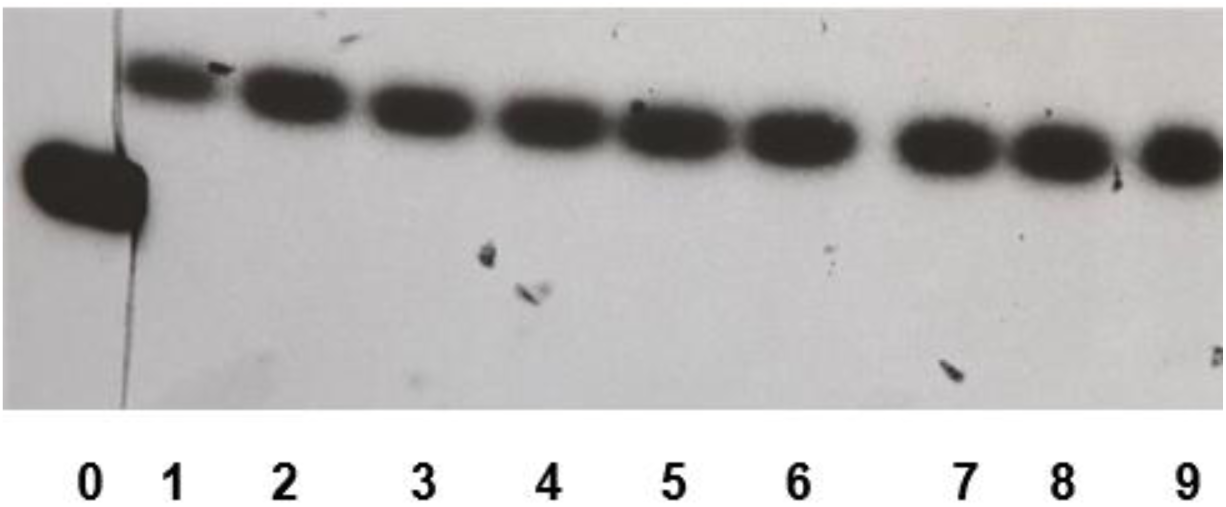
Modification of p(T)_15_ with [α-^32^P]o-GTP: **Lane 0:** 5′ end-labelled p(T)_15_ shown for reference. **Lanes 1 – 9**: modified oligo p(T)_15_. Aliquots were removed at various times over 24 hrs. and prepared for gel analysis. The enzyme was pre-incubated with the p(T)_15_ prior to o-GTP addition

The results of a corollary experiment are shown in Fig. 3b. In this experiment p(T)_15_ modification was done using [α-^32^P]o-GTP; 5′ end-labeled p(T)_15_ is shown as a reference to show the migration of unmodified substrate. An aliquot of the reaction in lane 9 (representing 24 hours) was also subjected to TLC and autoradiographed and yielding the same results as shown in Fig. 4; the oligo remained at the origin. The results shown in Figs. 3a, 3b, and 4 show clearly that the o-GTP is used as a substrate by TdT and able to be added to the 3′ end of the DNA substrate, and that aldehyde groups are present at the 3′ end.

**Fig. 4.**
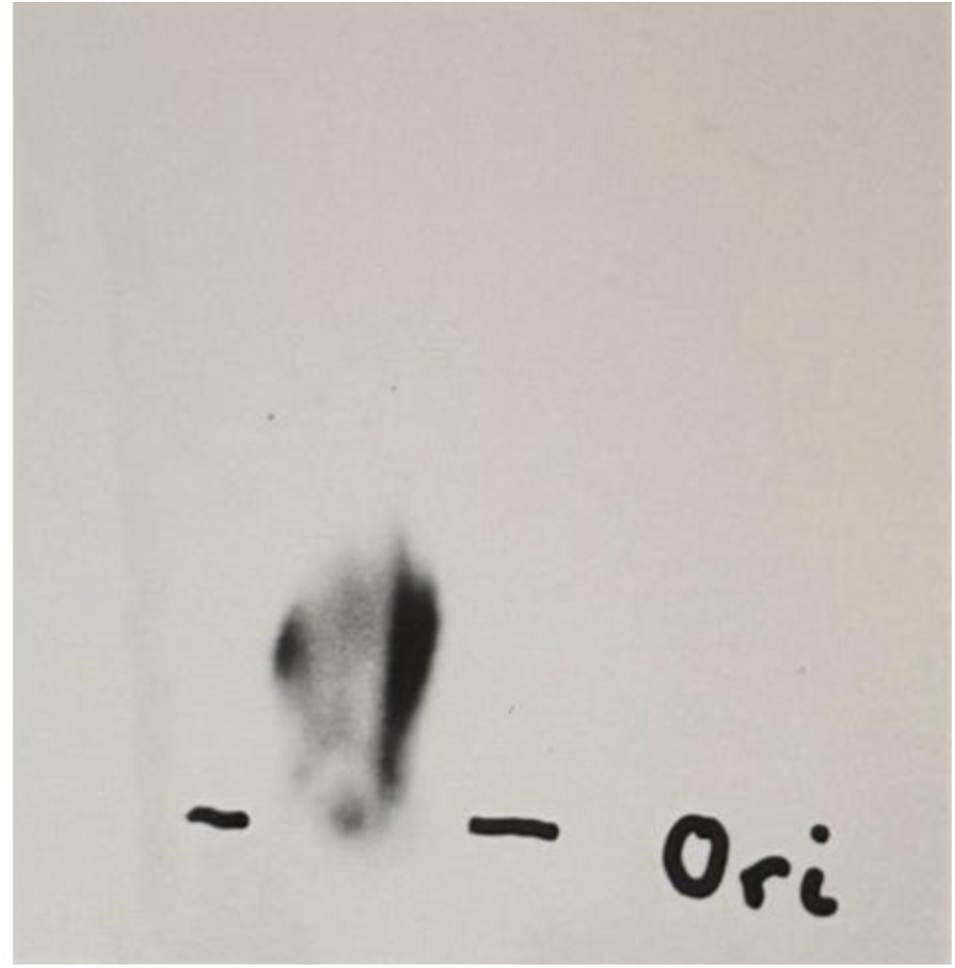
An autoradiograph of a thin layer chromatograph of 5′ end-labeled p(T)_15_ modified at the 3′ end with a 2′, 3′ *cis*-dialdehyde from o-GTP

Modification of DNA with aldehyde groups on the 3′ end has significance in that such a modification can permit covalent binding of the modified oligo to a sold-phase support for DNA synthesis or gene assembly, mutational studies, the synthesis of XNA, or for use in affinity chromatography to capture proteins that bind to DNA. These modified oligos may also be used as affinity labeling agents for proteins that interact with the 3′ end of DNA as demonstrated and discussed below. A diagram of the modified oligonucleotide is shown in Fig. 5.

**Fig. 5.**
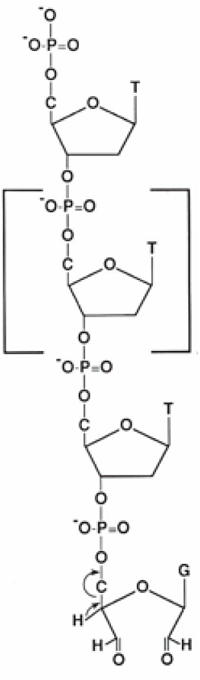
Diagram showing the *cis*-dialdehyde NMP on the 3′ end of ssDNA

The order of addition of the two substrates, DNA and o-NTP, is important. In all experiments with some exceptions (discussed below) the DNA substrate was incubated with TdT in the reaction mixture at least 10 minutes prior to addition the o-NTP.

#### (b) Demonstration of the utility of o-NTPs as possible affinity labeling agents

Very few reports appear in the literature where periodate-oxidized nucleotides have been used as affinity labels for DNA polymerizing enzymes. Perhaps this is because Srivastava et al. [11] investigated the use of o-ATP as an affinity agent against avian myeloblastosis virus reverse transcriptase and found that it bound reversibly to cysteine residues and concluded that o-ATP (or other o-NTP’s) may not be useful for selectively labeling lysine residues present at the substrate binding site of DNA polymerizing enzymes. We demonstrate that periodate-oxidized analogs of all four naturally occurring NTPs may be used as substrates by TdT to modify the 3′ end of oligonucleotide substrates which may then be used as affinity labeling agents.

If a compound is to be utilized as an affinity labeling agent, it is important to show that its binding to the protein is hindered by the presence of the physiological substrate which in this case is dGTP. As can be seen in Fig. 6, in the absence of dGTP (panel A) both the α and β subunits of TdT were labeled. In the presence of dGTP (Panel B), however, both subunits were labeled only minimally. This strongly suggests that o-GTP and dGTP compete for the same binding site.

**Figure 6.**
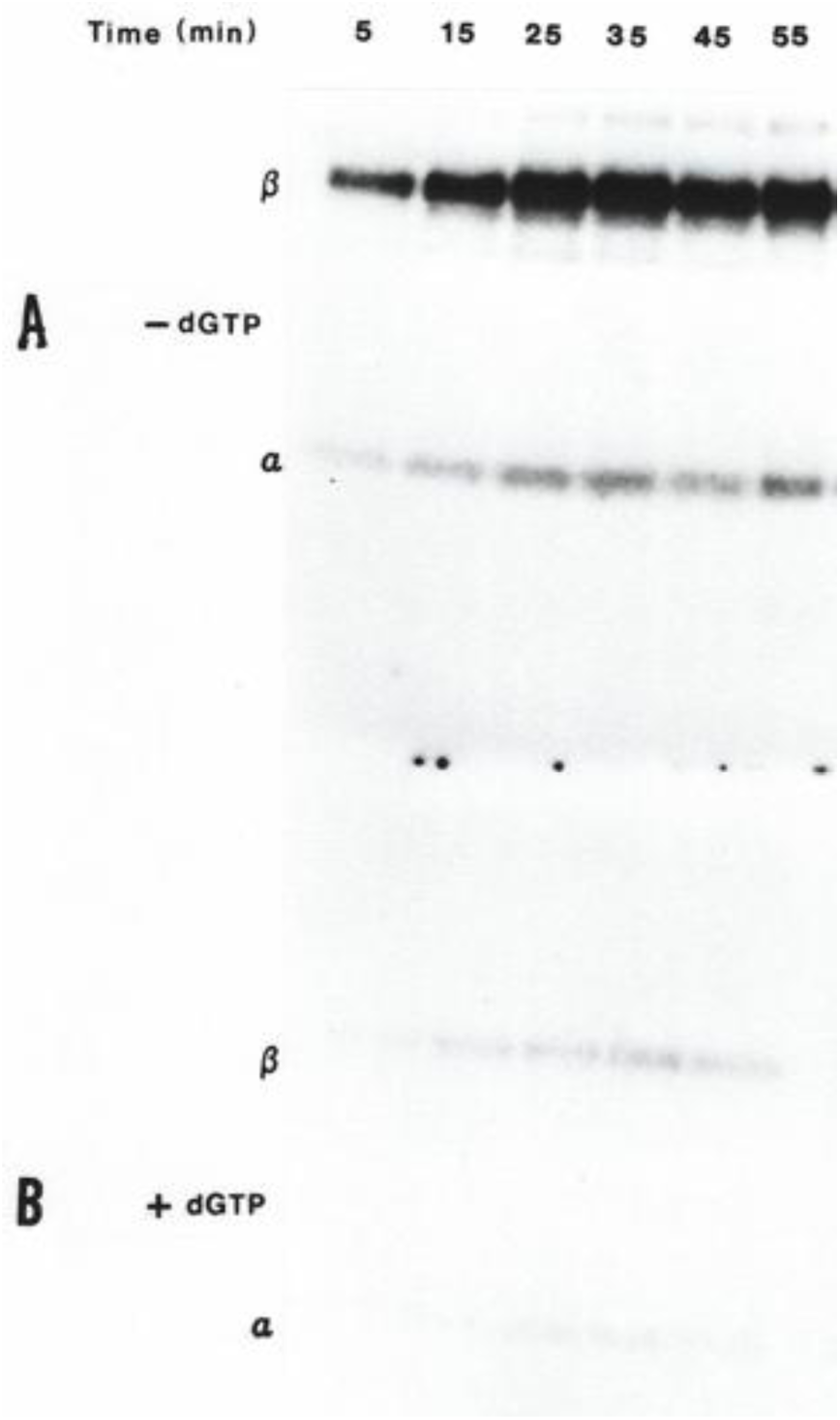
Gel analysis of protection against labeling TdT with o-GTP. Panel A represents TdT labeled in the absence of dGTP. Panel B shows protection by dGTP against labeling. All samples were treated with NaBH_4_ for 30 minutes prior to loading on the gel

Since TdT can also use oligonucleotides as a substrate, it is important to examine the effect of the oligo substrate on o-GTP binding. As can be seen in Fig.7 (Panel A), in the absence of the DNA substrate o-GTP binds to TdT as both the α and β subunits are labeled. When TdT is first incubated with the oligo substrate, o-GTP still binds as both the α and β subunits are labelled (Panel B). However, as shown, there is a change in the migration behavior of both the α and β subunits to yield the α′ and β′ subunits. Focusing on the β′ subunit, the change in migration is consistent with the 5′ end-labelled p(T)_15_ becoming covalently bound to the enzyme; presumably at the site where TdT interacts with the 3′ end of the oligo.

**Figure 7.**
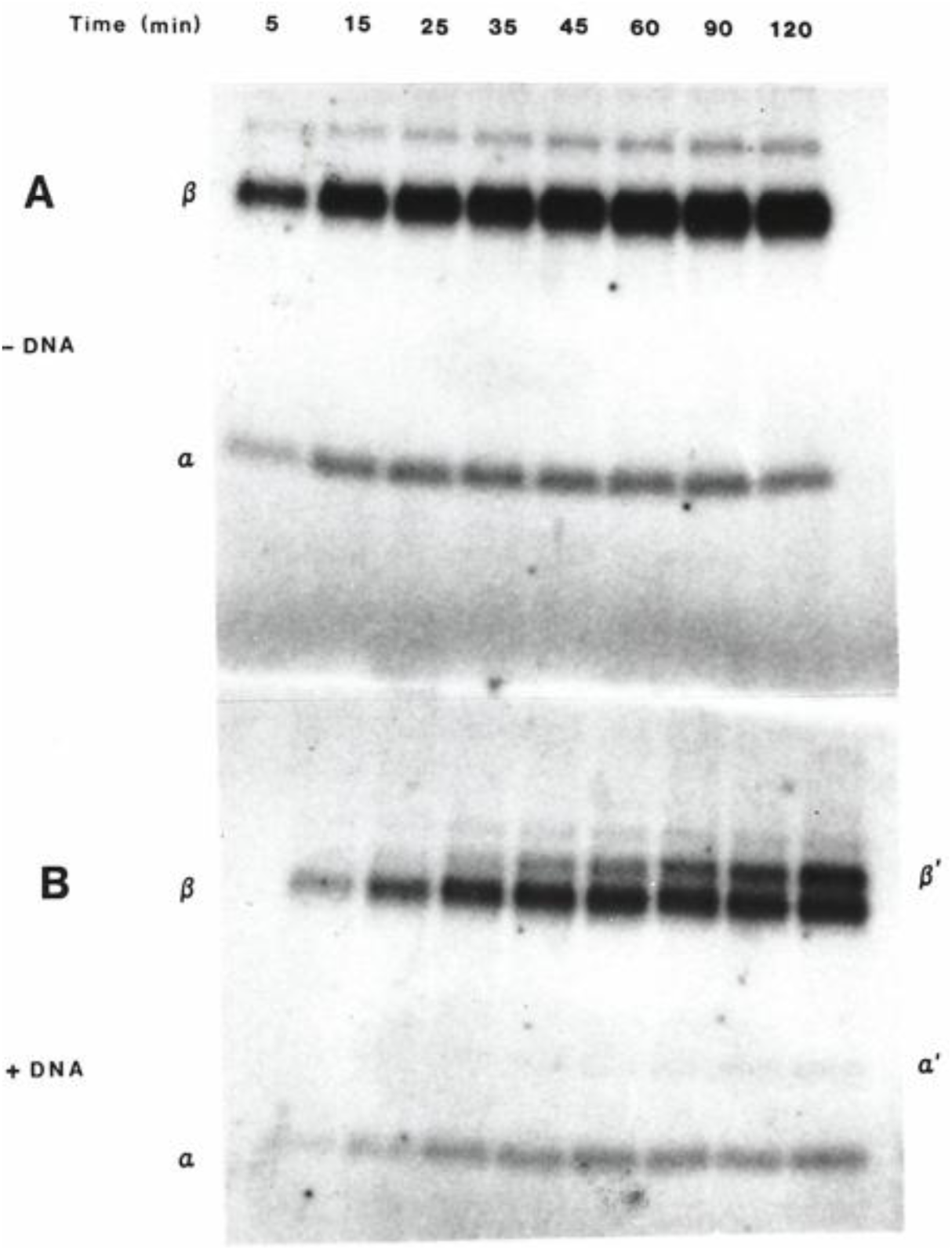
Labeling of TdT with 5′ end-labelled p(T)_15_ and o-GTP in the absence (Panel A) or presence (Panel B) of the oligo substrate. All samples were treated with NaBH_4_ for 30 minutes prior to loading on the gel

This experiment was repeated and the enzyme was then subjected to CNBr digestion, and the fragments were separated by SDS-PAGE. As can be seen in Fig. 8 in the absence of either oligo or dGTP several fragments are labeled (Control Lane) suggesting that o-GTP may also be binding non-specifically to sites other than the nucleotide binding site; possibly the DNA binding site. In the presence of dGTP, the enzyme is protected from o-GTP labeling. This is consistent with observations shown in Panel B of Fig. 6. In the presence of the oligo substrate o-GTP labels only one site on TdT (Fig. 8, +ssDNA lane). A possible explanation is that the oligo protects against labeling the DNA-binding site directly, or by effecting a conformational change such that the other sites are not amenable to labeling.

**Figure 8.**
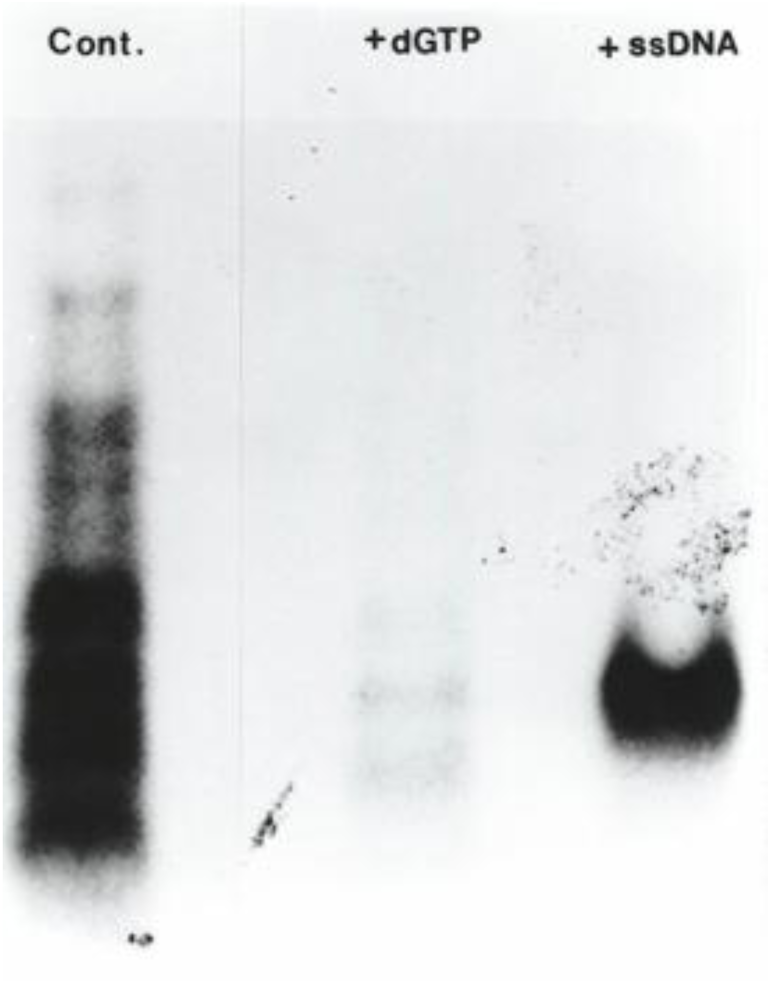
SDS-gel electrophoresis of CNBr fragments of TdT labeled with 5′ end-labelled p(T)_15_ in the presence of dGTP or ssDNA

Sequencing the fragment from the +ssDNA lane yielded **TGGFRRGKKIGHDVD** a sequence that shares sequence homology with several RNA and DNA polymerases [12, 13]. This sequence was also identified by Evans and Coleman [14] using the photo affinity label 8-azido ATP. This latter reagent contained a reactive azido group on the nitrogenous base while o-GTP contains the reactive aldehyde groups on the sugar moiety of the nucleotide. The fact that both reagents permit identification of a peptide from the same region of TdT suggests that the C-terminal region of the β polypeptide may interact with both the sugar and nitrogenous base of nucleotides.

The sequence reported here shares strong sequence similarity with polymerase β [15, 16]. Site-directed mutagenesis studies on amino acid residues in this region of polymerase β have implicated Arg-183 [17] and Asp-190 and 192 [18] as being involved in primer recognition and binding, respectively. Site-directed mutagenesis investigations on human TdT by Yang et al. [19] implicate the two Asp residues in this sequence as being part of the TdT active site. The ability to covalently attach an oligo modified at the 3′ end with one or two aldehyde groups to TdT after addition of NaBH_4_ suggests that this procedure may provide the basis for a novel affinity label which may be useful for studying of other proteins that interact with the 3′ end of DNA and possibly RNA.

Utilization of other o-NTP’s as substrates was examined. The results are the shown in Fig. 9, Lanes 2 – 4.

**Fig. 9.**
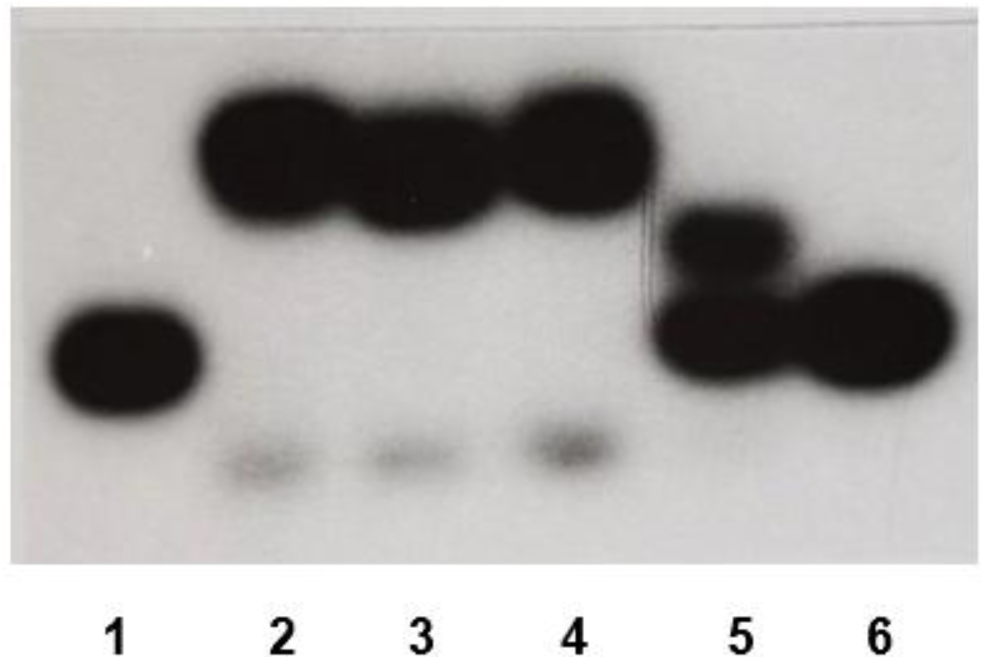
L**a**ne **1:** 5′ end-labelled p(T)_15_. In lanes 2 – 5 TdT was incubated with 5′ end-labelled p(T)_15_ before addition of the o-NTP substrate. Except for lane 5 which was heated, samples were not heat-treated prior to electrophoresis. Lanes 2 – 4 also show that even at 30°C (the reaction temperature) some of the primary product (upper band) is converted to the secondary product with a 3′ phosphoryl group (lower band). **Lane 2:** 5′ end-labelled p(T)_15_ modified using o-ATP. **Lane 3:** 5′ end-labelled p(T)_15_ modified using o-CTP. **Lane 4:** 5′ end-labelled p(T)_15_ modified using o-UTP. **Lane 5:** 5′ end-labelled p(T)_15_ modified using HO-ATP. Note the absence of a lower, secondary product band indicating that heating a 2′ 3′ *cis*-diol-modified oligo does not lead to formation of a 3′ phosphoryl group. **Lane 6:** TdT incubated with o-GTP prior to addition of 5′ end-labelled p(T)_15_. The lack of a primary product band suggests that o-NTP inhibits TdT if added prior to the DNA substrate

Terminal transferase was incubated 5 – 10 minutes at 30°C with 5′ end-labeled p(dT)_15_ *prior* to the addition of o-ATP, o-CTP, o-UTP, or HO-ATP. A change in the migration behavior of the radiolabeled oligo relative to p(dT)_15_ indicates that TdT is catalytically competent with all the o-NTP’s. In lane 6 TdT was incubated with unlabeled o-GTP for 15 minutes at 30°C *prior* to the addition of 5′ end-labeled p(dT)_15_. Apparently, brief exposure to the o-NTP before addition of the oligo substrate abolishes catalytic competence as evidenced by the lack of a change in the migration behavior of the oligo substrate in this lane. Presumably, addition of the o-NTP before adding the oligo substrate allows it to bind non-specifically to the DNA-binding site preventing DNA binding. Addition of the oligo substrate first protects against the non-specific binding of o-NTP to the DNA binding site so that it can bind only to the NTP binding site. Identical results were obtained when TdT was incubated with o-ATP, o-UTP or o-CTP prior to addition of the DNA substrate. Results shown here stand in contrast to those of Beabealashvilli et al. [20] and Chidgeavadze et al. [21] who examined several nucleotide analogues for their ability to inhibit and/or serve as substrates for various DNA polymerases of both prokaryotic and eukaryotic origin. These investigators provided data suggesting that TdT cannot utilize o-ATP as a substrate. A possible explanation for the apparent discrepancy between their results and those reported here may be that those investigators incubated TdT with o-ATP for a short period of time prior to addition of the DNA substrate. These results also reveal that the o-NTP binds to lysine residues not reversibly to cysteine as suggested by Srivastava et al. [11].

Results reported here indicate that the order of addition of substrates is crucial; the DNA substrate must be added before the o-NTP. Even short incubation times of TdT with the o-NTP prior to addition of the DNA substrate leads to inhibition of TdT and thus no o-NTP incorporation into DNA is observed.

#### (c) Demonstration of 3′-phopsphate addition

Studies *in vitro* have shown that TdT can modify the 3′ end of oligonucleotides with a phosphoryl group using specially modified deoxynucleoside triphosphates [22, 23]. Camden et al. [24] have shown that a phosphoryl group can be added to the 3′ end of DNA or RNA oligonucleotides by a synthetic oligonucleotide 3′ kinase1 deoxyribozyme using PPP-5′-RNA as the phosphoryl donor. Flamme et al. [25] have shown that TdT may be used to generate DNAs with a 3′-phosphoryl group using specially synthesized nucleotides which contain a modified phosphoryl group on the 3′ end. When the modification is removed, a DNA with a 3′-phosphoryl group is left behind. Here we show that oligonucleotides may also be phosphorylated at their 3′-ends using TdT and o-NTPs followed by heating in the presence of amino groups on the enzyme.

In the experiments depicted in Figs. 3a and 3b samples were not heated prior to loading onto the gel for electrophoresis. When samples were heated before electrophoresis, the initial upper product band disappeared, but a lower secondary product band appeared just below the p(T)_15_ substrate band (Fig. 10). Heating the modified oligo for increasing time resulted in more primary product being converted to a secondary product.

**Fig. 10.**
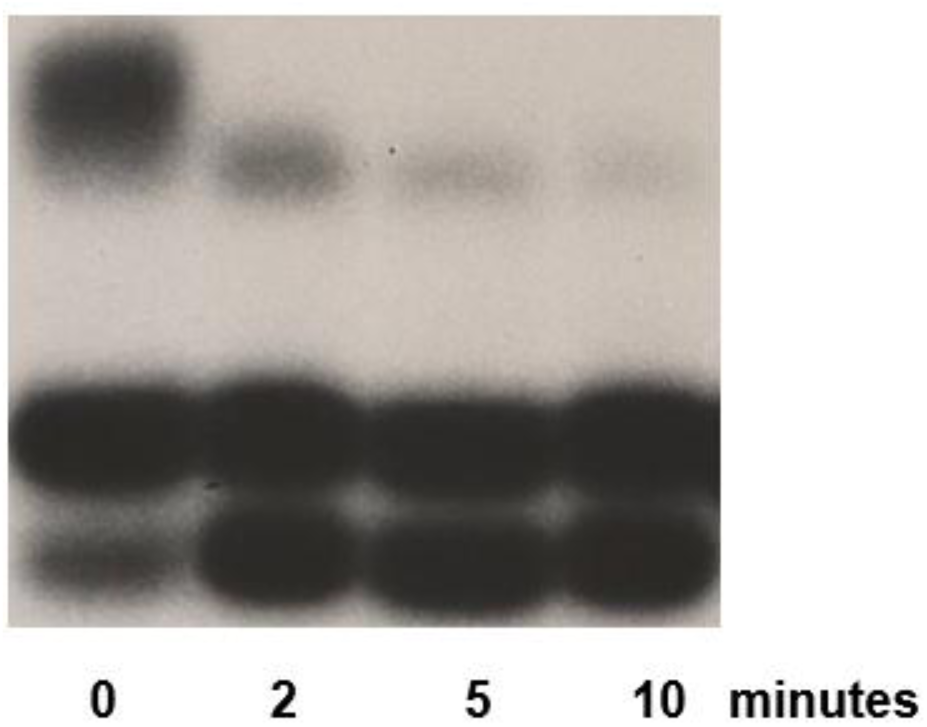
Illustrates the effect of heating the 2′, 3′ *cis*-dialdehyde modified 5′ end-labelled p(T)_15_ at 95°C for varying amounts of time. The primary product (upper band) disappears with increased time of heating from 0 minutes to 10 minutes, and a secondary product (lowest band), representing the oligonucleotide with a 3′-phosphoryl group, increases. The observation of a secondary product band at 0 minutes indicates that even at 30°C (the reaction temperature) some primary product is converted to secondary product

An aliquot of p(T)_15_ whose 3′ end was modified with [α-^32^P]o-GTP was subjected to TLC and autoradiographed (Fig. 11). Consistent with the oligo possessing aldehyde groups at its 3′ end, the unheated modified p(T)_15_ remained at the origin (Lane 2). Heating effected a change in the migration of p(T)_15_ such that the oligo did not remain at the origin yet retained the α-^32^P (Lane 1). The results are consistent with the oligo having aldehyde groups at the 3′ end before heating, then losing those groups upon heating.

**Fig. 11.**
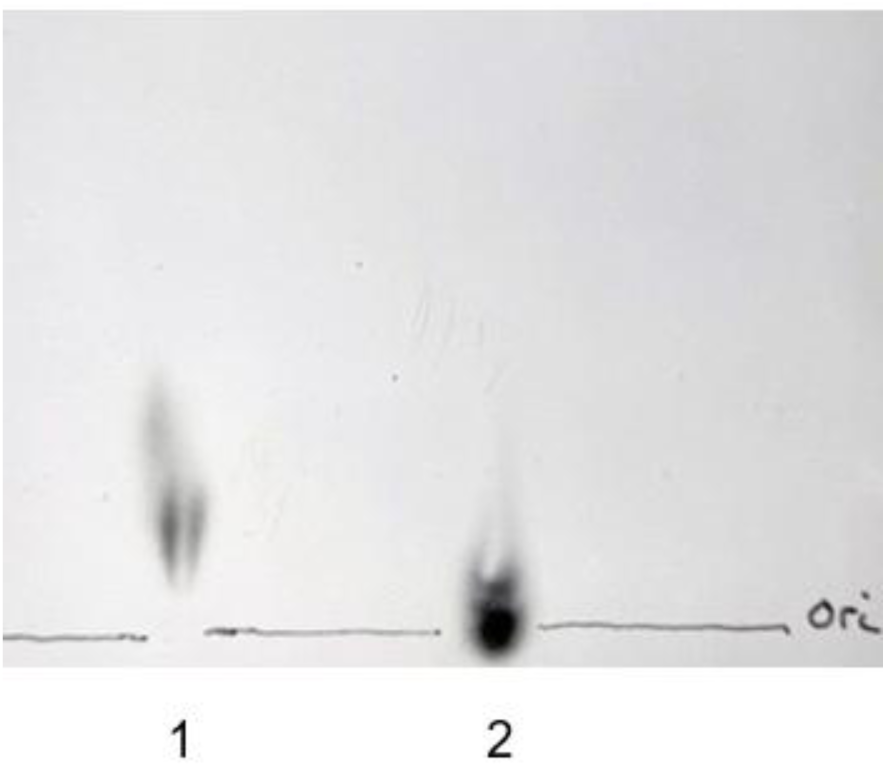
Autoradiograph of a thin layer chromatograph of p(T)_15_ modified at the 3′-end. **Lane 2:** modified with [α-^32^P]o-GMP *cis*-dialdehyde before heat-treated. **Lane 1:** shows the same modified p(T)_15_ after heat-treated

The results of a complementary experiment, shown in Fig. 12, also revealed that the ^32^P which remains with the heat-treated oligo is the α-phosphate of the o-NTP substrate.

**Fig. 12.**
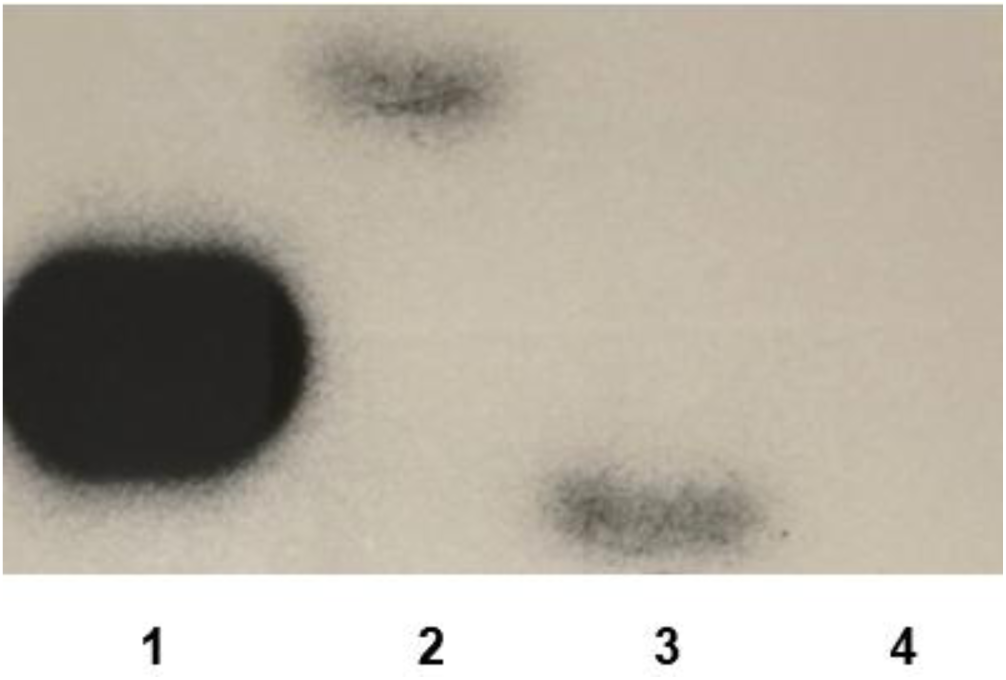
L**a**ne **1:** 5′ end-labelled p(T)_15_. **Lane 2:** p(T)_15_ labelled on the 3′ end with [α-^32^P]o-GTP 2′, 3′ *cis*-dialdehyde. **Lane 3:** The product in lane 2 heat-treated for 10 minutes; p(T)_15_ with a 3′-phosphoryl group. **Lane 4:** The product band in lane 3, note that the labeled phosphoryl group is removed after treatment with alkaline phosphatase

Lane 1 is unmodified 5′ end-labeled p(T)_15_ for reference. Lane 2 is p(T)_15_ modified with [α-^32^P]o-GTP 2′, 3′ *cis*-dialdehyde at its 3′ end. Lane 3 is an aliquot from the sample in Lane 2 that has been heat-treated. Lane 4 is an aliquot from the sample in Lane 3 that was treated with alkaline phosphatase which should remove the 3′-phosphate group. The result shown in Lane 4 clearly indicates that the phosphate group remaining attached to the 3′ end after heating (Lane 3) is the α-phosphate.

A model depicting the reaction upon heating is shown in Fig. 13. The uppermost band, the primary product (Lane 2, Fig. 12), is represented on the left in Fig. 13. When heated in the presence of amine groups (e.g., amino groups in the protein), it undergoes a β-elimination reaction generating a secondary product (Lane 3, Fig. 12, and the right side of Fig. 13). The oligo has the same nucleotide length as p(dT)_15_, but with a 3′-PO_4_^2-^, replacing the 3′-OH. The added -PO_4_^2-^ gives the oligonucleotide an additional -2 charge, causing it to migrate a little further toward the anode during electrophoresis and thus generate a secondary product band just below the p(T)_15_ band. Figure 13 also shows the unsaturated *cis*-dialdehyde nucleotide in a dihydroxymorpholino linkage with an amino group on the enzyme. This is not to imply that only this type of linkage is capable of β-elimination. The elimination reaction can also be promoted by formation of a Schiff’s base between an amino group and the C3′ aldehyde group of the o-NTP [27].

**Fig. 13.**
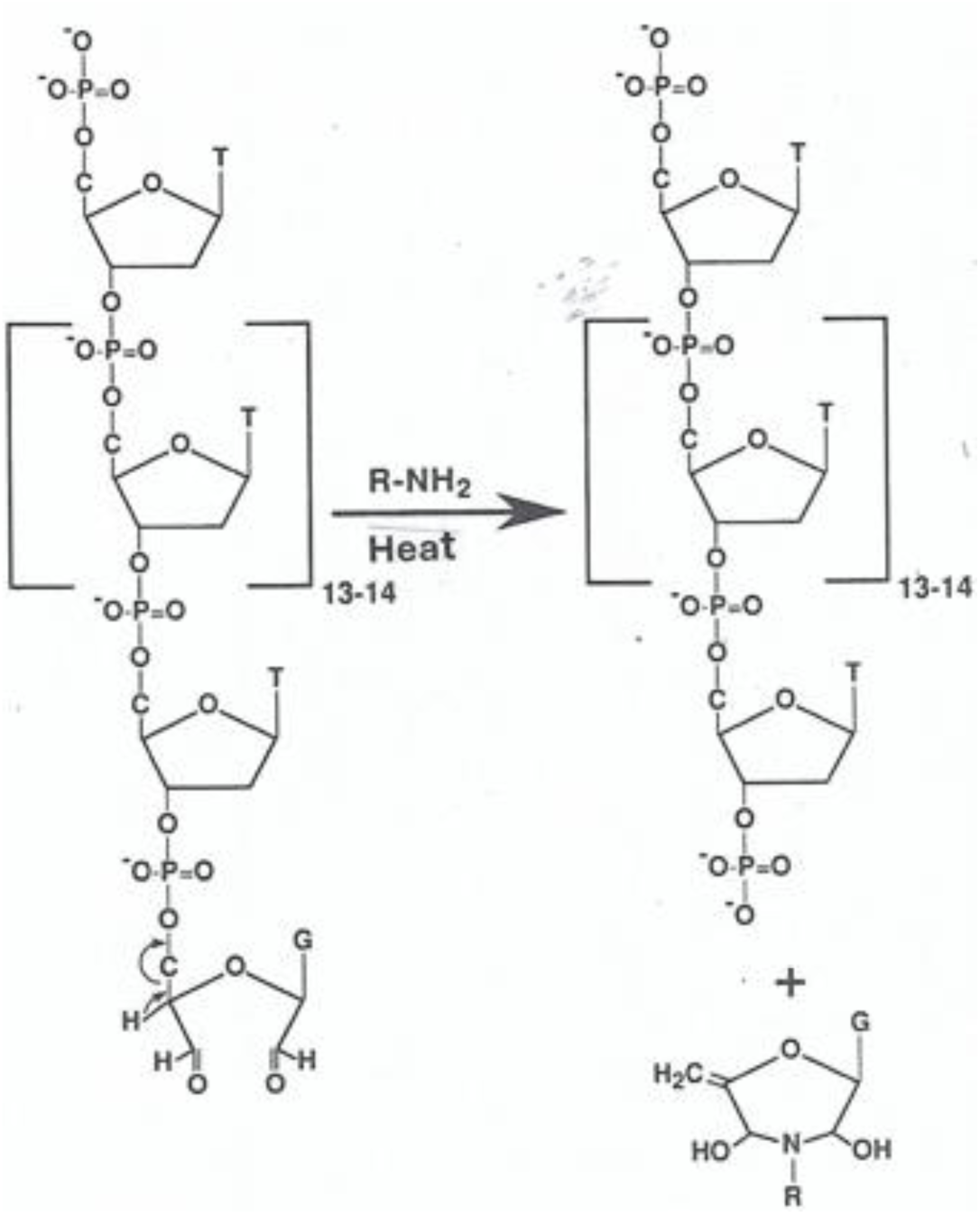
is a model of the formation of an oligonucleotide modified with a phosphate group at its 3′ end by heating the oligonucleotide modified with a 2′, 3′ *cis* dialdehyde

These experiments show clearly that a 3′-phosphoryl group can easily and efficiently be added to DNA or RNA oligonucleotides by using TdT to first catalyze the addition of a o-NMP to the 3′ end then heating for several minutes. Oligonucleotides possessing a 3′-phosphoryl group may then be ligated to other oligonucleotides that have a 5′-OH as described by Zhelkovsky and McReynolds [1]. This may prove useful in the synthesis of long strands of synthetic RNA, the synthesis of single-stranded RNA/DNA hybrids, or the synthesis of long-stranded XNAs.

#### (d) Demonstration of 2′, 3′-cis diol NMP addition

Terminal transferase is also capable of using HO-NTP as a substrate with 5′ end-labeled p(T)_15_ as shown in Fig. 9. Lane 5 shows a product band just above the DNA substrate band. Note that the migration of an oligo modified with *cis*-diols differs significantly from an oligo modified with *cis*-dialdehydes (Lanes 2 – 4). The observation that a significant amount of the DNA substrate remains unreacted suggests that the *cis*-diol substrate is utilized less efficiently than the *cis*-dialdehyde substrate. When heated, diols do not undergo a β-elimination [26] consequently, there is no secondary product band below the starting material.

Periodate-oxidized nucleotides possess two reactive aldehyde groups on the 2′ and 3′ carbons. Perturbation of the ribose ring with periodate allows more freedom of rotation around carbons C2′ and C3′ and around the C4′ – O – C1′ bonds. Extensive structural studies on o-ATP reveal that periodate-oxidized nucleotides react with proteins like free aldehydes, however, in solution the presence of free aldehyde groups is not apparent [4, 27]. Both ^1^H-NMR and ^13^C-NMR studies suggest that the aldehyde groups are hydrated leading to the formation of a hemiacetal between C3′ and C2′ forming a six-membered ring structure. The -OH groups on carbons C2′ and C3′ may then adopt any of four different axial or equatorial conformations with respect to the ring [4]. In each case the hemiacetal can mimic, to some degree, the ribose ring decreasing the freedom of rotation about bonds. When reacting with a protein, all or only one of the conformations may interact favorably enough to permit product formation.

Unlike the *cis*-dialdehyde, a *cis*-diol substrate cannot form a hemiacetal [4] thus there is much more freedom of rotation around bonds suggesting that the *cis*-diol may not react as favorably as the *cis*-dialdehyde substrate. This may explain why a significant amount of the DNA substrate appears to remain unreacted as shown in Fig. 9. It could be argued that the change from a planar sp^2^ configuration about the C2′ and C3′ aldehyde carbons of an o-NTP to a tetrahedral sp^3^ configuration upon reduction to hydroxyls [4] in an HO-NTP effects a decrease in the affinity between the HO-NTP and TdT compared to o-NTP.

### CONCLUSIONS

Terminal transferase can efficiently use periodate-oxidized nucleotides as substrates to modify the 3′-end of a nucleic acid with *cis*-dialdehydes. In the case of TdT the dialdehyde-modified nucleic acid may then effectively be used as an affinity label to identify amino acid residues in or near the active site of proteins that interact with the 3′-end of DNA.

Oligonucleotides modified with a 3′-phosphoryl group may be generated by simply heating the dialdehyde-modified oligonucleotide, the dialdehyde is induced to undergo a β elimination reaction leaving behind its α-phosphate group on the 3′-end.

Terminal transferase can use periodate-oxidized nucleotides that have been reduced by NaBH_4_ to modify the 3′-end of oligonucleotides with *cis*-diols. The use of these reduced nucleotides is not as efficient as the dialdehydes due to increased rotations around certain bonds in the nucleotides and the fact that, unlike their dialdehyde counterparts, they do not form hemiacetals.

Thus, TdT may be used in a simple, straightforward procedure to efficiently modify the 3′-end of oligonucleotides with dialdehydes, diols, or phosphoryl groups. This report expands the many uses of TdT as a tool in the molecular biology toolkit.

## Competing Interests

There were no competing interests of the authors.

## Author Contributions

RSA Conceptualization, experimental set up, data analysis, writing and editing KLB Conceptualization, editing, advising and Principal Investigator

## Acknowledgements

Support for this work came from NIH grant GM25530 and the Baylor College of Medicine Department of Biochemistry RSA is indebted to Drs. Roseann Cappel and Nihimat Morjana for many helpful discussions and advice.

To Shaveen De Mel for assisting with the figures.

## Data Availability

Individuals may request additional data by contacting RSA directly.

## Notes

### Competing Interest Statement

The authors have declared no competing interest.

